# Determining the Statistical Significance of the Difference Between Arbitrary Curves: A Spreadsheet Method

**DOI:** 10.1101/2023.04.06.535947

**Authors:** Kalina Hristova, William C. Wimley

## Abstract

We present a simple, spreadsheet-based method to determine the statistical significance of the difference between any two arbitrary curves. This modified Chi-squared method addresses two scenarios: A single measurement at each point with known standard deviation, or multiple measurements at each point averaged to produce a mean and standard error. The method includes an essential correction for the deviation from normality in measurements with small sample size, which are typical in biomedical sciences. Statistical significance is determined without regard to the functionality of the curves, or the signs of the differences. Numerical simulations are used to validate the procedure. Example experimental data are used to demonstrate its application. An Excel spreadsheet is provided for performing the calculations for either scenario.

## INTRODUCTION

Biological phenomena are often characterized by the measurement of curves portraying a dependent experimental output as a function of an independent parameter such as concentration, dilution factor, or time. Some biological phenomena are well described by explicit, physically meaningful functions, and thus can be reduced by curve fitting to a set of physically meaningful fit parameters, with associated uncertainties, that can be compared statistically. But many biological phenomena obey complex functions that cannot be described with mathematically simple functions that have physically meaningful parameters. Thus, determining the statistical significance of the difference between curves is a commonly faced problem that does not have a widely known and easily implemented solution.

The problem of comparing arbitrary curves is one of testing the null hypothesis that the two curves, such as the simulated data in **Fig. 1**, were sampled from the same parent population, at each value of X, but without any assumptions about how the curve values change with X. Ultimately, comparing arbitrary curves requires a comparison of the total sum of squared differences between the curves to the distribution of differences expected from random sampling when the null hypothesis is true. This goal is not accomplished by two-factor ANOVA or by any other commonly used or widely known method in biostatistics.

**Figure 1.**
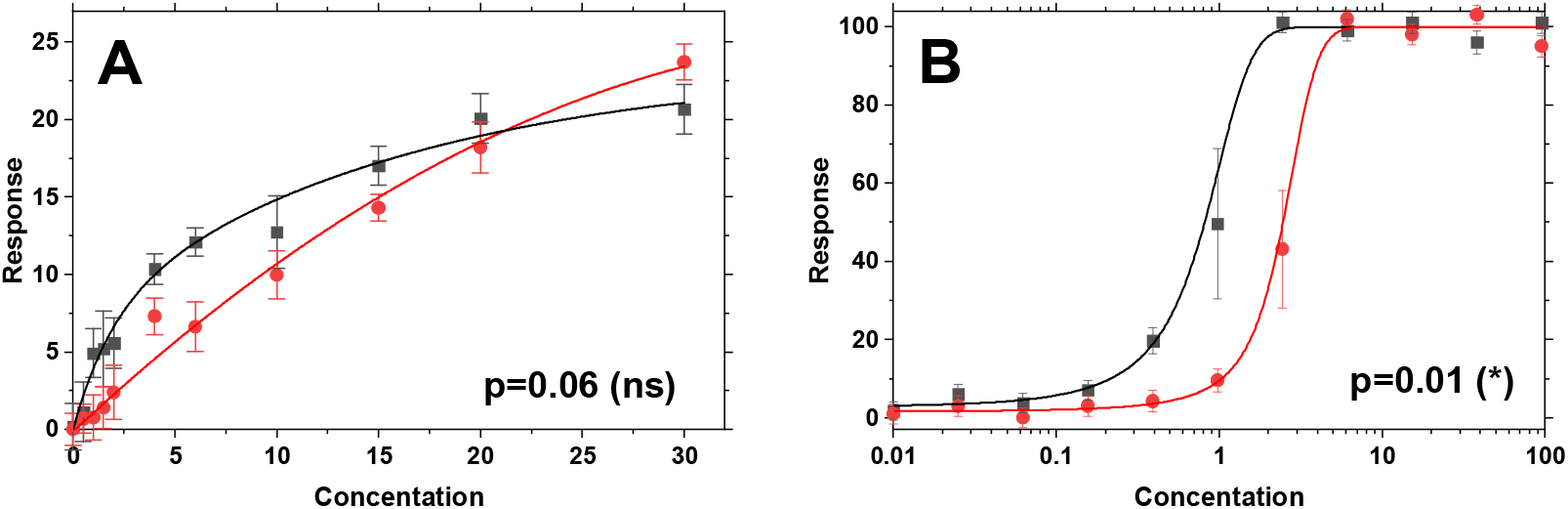
Simulated pairs of curves containing small systematic differences. The uncertainties at each point are obtained by randomly sampling from a Gaussian distribution with N=4. **A**. Curves of a type normally plotted on a linear X-axis. In this example, uncertainties are constant (independent of Y) and are shown as standard errors calculated individually from randomly sampled data for each point. **B**. Curves of a type typically plotted on a log scale X-axis. In this case uncertainties scale with the slope of the curve. Using the modified Chi-squared method described here, any pair of arbitrary curves with known uncertainties, such as these, can be compared to produce a p-value for the null hypothesis that both curves were sampled from the same parent population at each value of X. The p-values for the two curves, determined with the technique described here, are shown.

Here we present a simple spreadsheet-based method of determining the statistical significance of the overall difference between any pair of arbitrary curves. This method is based on the commonly used Chi-squared test for goodness of fit [1] but with an essential correction for the deviation from normality that arises with small to moderate sample sizes, which are nearly universal in biomedical research. This method enables a user to calculate an overall p-value for the null hypothesis that, at every value of X, the measurements for two curves were sampled from the same parent population. This is achieved with a false positive rate (at least one Tyle I error over all comparisons) equal to α, the chosen cutoff for statistical significance, typically 0.05. The method we describe also provides a post hoc test for calculation of p-values for the individual pairs of points at each value of X, with a correction for multiple comparisons, so the user can determine which points contribute most to the overall statistical significance.

## METHODS

### Chi-squared comparison of two means

The Chi-squared (χ^2^) probability distribution was first described in 1900 by Karl Pearson to compare categorical data [2], but it has also found many applications in continuous data, especially in regression and curve fitting [1]. In goodness of fit tests, χ^2^ takes the general form:

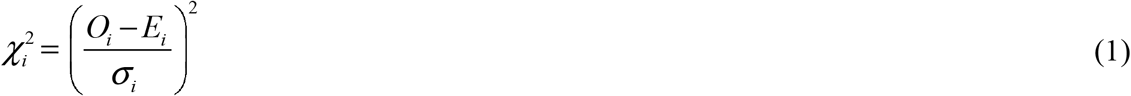

where O_i_ is the observed value of point i, E_i_ is the expected value of point i based on the fit curve, and *σ*_i_ is the standard deviation of O_i_ in the Y-dimension at point i.

The utility of the χ^2^ formalism is that the individual 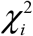 values can be summed over any number of points to determine an overall statistic for the sum of the differences. This is accomplished without increasing the probability of false positive results that arises due to multiple comparisons. This sum of χ^2^ enables calculation of a probability, the p-value, for the null hypothesis that the differences were sampled from parent population with a mean of zero, i.e. that the differences between the curves arose only due to random sampling. Neither the signs of the differences nor the absolute values of the points influence the final results of this analysis, which are dependent only on the magnitude of the difference between points, relative to the uncertainty in the measurements. Here, we will show how to use this method to generate p-values for the pairwise comparison of any two experimentally determined curves, using two different scenarios.

### Scenario 1: One measurement at each X with known SD

To determine the statistical significance of the difference between curves, one must have a measure of uncertainty for each point. If each point in the curve has been measured in a single experiment, then one must have knowledge of the standard deviation *σ*_*i*_ in that measurement. Ideally, such knowledge should be quantitative and based on previous measurements using the same experimental technique. When comparing a pair of measurements, 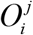, where j = dataset a or b, each determined from a measurement with an associated 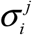, then 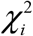 takes the form:

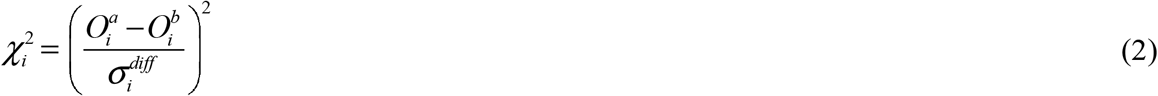

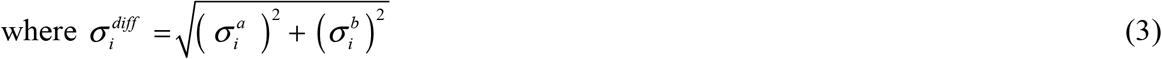

The standard deviation of the difference is calculated by propagation of errors [1].

*NOTE: In this scenario, the steps described next in Equations 4-9 are not applicable. The reader may skip to Equation 10. The* ***Supplemental Spreadsheet*** *includes this scenario in tabs indicated by “N=1 with SD”*.

### Scenario 2: N measurements at each X, giving mean and SE

The best case is to measure each point of each curve in N independent experiments, so that each point can be represented as a mean *O*_*i*_ and its measured standard error *SE*_*i*_ . When comparing a pair of such measured means, 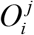, where j = dataset a or b, each determined from N^j^ independent measurements, 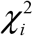 takes the form

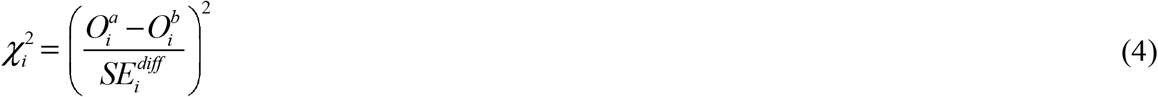

In this equation, 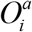 and 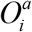 are the two means and 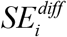 is the standard error of the difference, defined by

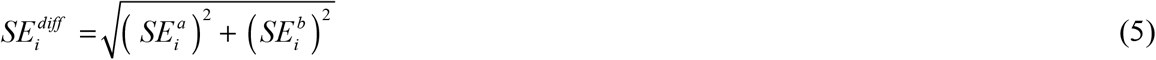

where 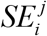 is the standard error of each measurement, calculated from the measured standard deviation,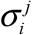

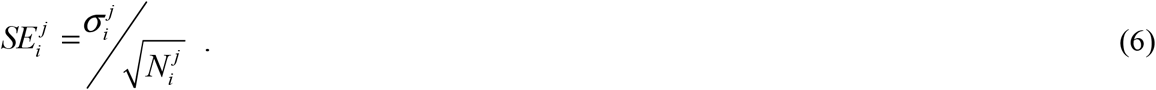

### Using the t-distribution to correct for small N

The χ^2^ formalism just described for a measurement with SE assumes that the parent probability distributions in Y around each point follow a Gaussian/normal distribution shape. However, as Gossett (aka “Student”) first showed in 1908 [3], this is only true when the mean of each point is determined with large N. In biomedical sciences, this condition is rarely satisfied. Severe undersampling is typical. For N _≤_ 20 the deviations from normality are not trivial, and for N_≤_10, they are large and cannot be ignored. Gossett thus defined the t-distributions as the appropriate probability distributions to use for small N values. To correct for the effect of small N, while still using the χ^2^ formalism, we will determine a corrected χ^2^ for each pair of points using t-distributions, instead of directly calculating it using equation 4.

In this method, the t-distribution is used to calculate the corrected probability from the differences between each pair of points compared to the uncertainty, assuming the null hypothesis is true and the samples are normally distributed at large N. We note that the t-test probability does not assume that the SD of the two points are the same. Thus, the t-value and probability are equivalent to those from a Welch’s corrected, two-sample t-test, which does not assume that SDs are equal. This probability is then used to calculate an equivalent, thus corrected, value for 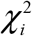.

To use the t-distribution to calculate probability, we first calculate t for each pair of points

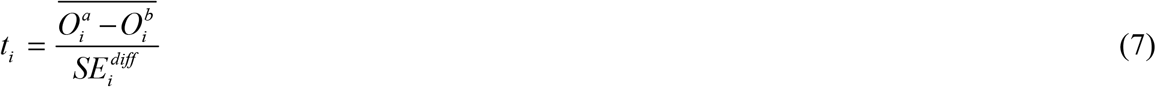

where the terms are defined above. With t and dF^t^ (degrees of freedom for t), the probability for each pair of means is determined using a t-distribution calculator.

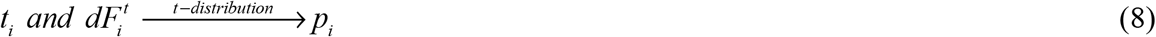

(We note that the calculators mentioned are built in to most spreadsheet/programming environments including Microsoft Excel, see below). In turn, this probability is used to calculate a corrected Chi-squared with an inverse Chi-squared calculator using dF=1 for the pair of points.

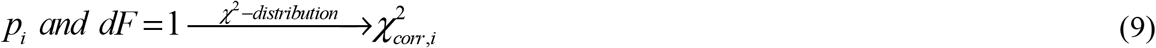

### Calculating the overall statistical significance

To calculate the overall p-value of the difference between curves, over all pairs of points taken together, the corrected χ^2^ values are summed

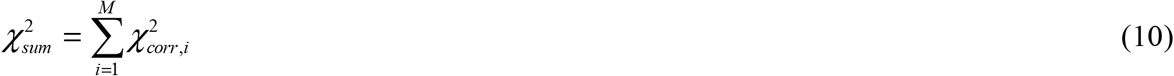

and then the overall p-value for the comparison of two curves is determined using a χ^2^ calculator using dF=M, where M is the number of pairs of points.

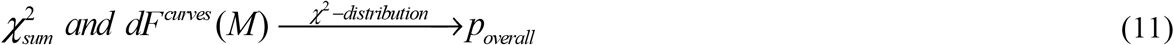

### The null hypothesis and p-value for comparison of curves

The null hypothesis for a comparison of two arbitrary curves is that each pair of measurements, 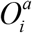 and 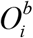, collected at the same value of X, were sampled from the same parent population at that point. There is no assumption about how the parent populations differ from one another at different X values, thus the differences between any pair of curves with arbitrary shape can be assessed.

### Determination of SE values

An accurate measurement of uncertainty based on multiple independent experiments is the best case for calculating accurate p-values using this method. When comparing two curves, ideally one should use uncertainties that are individually determined for each point (scenario 2). If uncertainties are expected to be equal across the range of X, averaged SE values can be used as long as they are experimentally determined using independent measurements. In simulations like those described below, we found that there is no systematic difference between individually measured and averaged uncertainties, as long as they are not systematically different across the curve.

### Degrees of freedom for t-distribution

Typically, the individual means being compared will both be measured with the same N points, such that dF^t^ = 2N-2. When the two means are measured with different number of points, N_larger_ and N_smaller_, the smaller N dominates the deviation from normality. In this case, the best estimate for dF^t^ is dF^t^ = 2N_smaller_ – 2.

### Spreadsheet method

We show next how the overall p-value for the comparison of two arbitrary curves can easily be determined using a Microsoft Excel spreadsheet. A spreadsheet with these functions coded as described below is included in **Supplemental Information**. Options for the two scenarios described above are included in the spreadsheet: Scenario 1: N=1 measurement at each X, with a known SD (labelled “N=1 with SD”) or Scenario 2: N > 1 measurements at each X which are used to calculate mean and SE (labelled “N>1, mean and SE”). These functions are easily recreated in MatLab, Python, or other programming environments using built-in functions.

### To calculate the individual χ2 values in Excel when N=1 and SD is known

Column designations match the **Supplementary Spreadsheet** labelled “N=1 with SD”

Col A: Enter *X*_*i*_, the X-values for the two curves, a and b

Col B: Enter 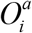 (Measured value of Y for point i, Curve a)

Col C: Enter 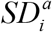 (SD of point i, Curve a)

Col D: Enter 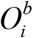 (Curve b)

Col E: Enter 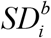 (Curve b)

Col G: Calculate absolute value of difference between the means of the ith pair of points with [=ABS(Di-Bi)]

[Note: characters within brackets refer to excel formulas with the column designations in the supplementary spreadsheet]

Col H: Calculate standard deviation of the difference between ith pair of points, with [=SQRT(Ci^2+Ei^2)]

Col I: Calculate the individual Chi-squared values by [=(Gi/Hi)^2]

### To calculate mean and SE when N>1 in EXCEL

Enter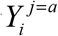 raw measurements for curve *a* in rows, one row per X-value

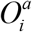- Calculate mean of point i in curve j=*a* with [=AVERAGE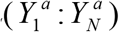] where N is 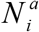

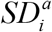 - Calculate SD of point i in curve j=a with [=STD.S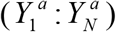] where N is 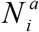

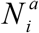 - Calculate number of measurements for each point i in curve j=a with [=COUNT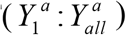]

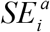 - Calculate SE of point i in curve j=a with [=STD.S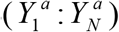/SQRT(N)] where N is 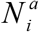

Repeat above steps for curve b to get 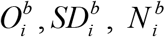 and 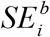

Paste these values into the worksheet for Chi-squared calculation.

### To calculate the individual Chi-squared values in Excel with N>1, mean and SE

Column designations match the **Supplementary Spreadsheet** labelled “N>1 with mean and SE”

Col A: Enter *X*_*i*_, the X-values for the two curves, a and b

Col B: Enter 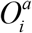 (Mean of Y for point i, Curve a)

Col C: Enter 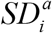 (SD of point i, Curve a) (SD is not used directly in the calculations)

Col D: Enter 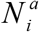 (Number of measurements for point i, Curve a)

Col E: Enter 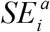 (SE of point i, Curve a)

Col F: Enter 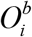 (Curve b)

Col G: Enter 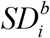 (Curve b)

Col H: Enter 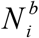 (Curve b)

Col I: Enter 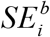 (Curve b)

Col K: Calculate absolute value of difference between the means of the ith pair of points with [=ABS(Fi-Bi)]

Col L: Calculate standard error of the difference between ith pair of points, with [=SQRT(Ei^2+Ii^2)]

Col M: Calculate t-value for ith pair of points with [=Ki/Li]

Col N: Determine N for dF calculation. If the number of data points is unequal, choose the smaller of the two N values, with [=IF(Hi<Di,Hi,Di)]

Col O: Calculate dF for t-calculation with [=2*Ni-2]

Col P: Calculate the probability from t and dF with [=T.DIST.2T(Mi,Oi)]

Col Q: Calculate corrected 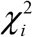 using the probability in Col P and dF=1 with [=CHISQ.INV.RT(Pi,1)]

### To calculate the overall p-value in Excel using χ^2^

The attached **Supplemental Spreadsheet** includes two scenarios for uncertainties:

Scenario 1: N=1 with SD from prior knowledge, indicated in the spreadsheet tab by “N=1 with SD”. In this scenario, “Col X” below refers to Col L and “Col Y” below refers to Col I.

Scenario 2: SE from multiple measurements, indicated in the spreadsheet tab by “N>1, mean and SE”. In this scenario, “Col X” below refers to Col T and “Col Y” refers to Col Q.

Col X, Cell 2: Calculate sum of 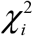 with [=SUM(Y3:Ym)] where m is the row containing the last pair of points

Col X, Cell 3: Determine dF for 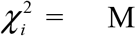, the number of pairs of points, with [=COUNT(Y3:Ym)]

Col X, Cell 4: Calculate overall p-value for the comparison of the two curves using the sum of χ^2^ and dF with [=1-CHISQ.DIST(X2,X3,TRUE)]

## RESULTS and DISCUSSION

### Validation by simulation

To validate this entire method, for both scenarios, including the correction for deviation from normality, and the choices for degrees of freedom, we performed numerical simulations of 6,000 pairs of curves. Curves were created by randomly sampling points from a Gaussian parent population to generate a simulated experiment that satisfies the null hypothesis that all points were sampled from the same parent population at each value of X. For each simulation, we determined the overall p-value using the modified χ^2^ method as above. We tested datasets with 1, 3 and 10 measurements per point (N) and with 3 or 10 points per curve (M). Because the simulated data are random, and satisfy the null hypothesis by definition, we can validate the modified Chi-squared method by showing that the frequency distributions of observed p-values and observed χ^2^ values obtained from the simulations match the expected frequencies. More importantly, the frequency of simulation outliers that give false positive results (i.e. Type I errors) can be compared to the critical right tail of the asymmetric χ2 distribution.

In the simulations shown in **Fig 2A**, we sampled points with N=3 from a Gaussian parent distribution. Simulations with 3 measurements per point are a worst case for the necessary correction for the deviation from normality that arises due to small N. In **Fig. 2C** we sampled points with N=10. We compared the simulated curves using the modified Chi-squared method, and accumulated 6,000 overall Chi-squared values and overall p-values. We binned the p-values into 6 bins; two 0.025 wide probability bins, each, on the left and right side of the distribution and the remaining 0.90 probability bin in the middle of the distribution. To test for outliers, we also measured the probability of observing p≤0.001. The results in **Fig. 2A&C** show that the p-value frequencies are correct when N=3 and when N=10 for bin sizes of 0.025 and 0.05, for both left and right tails, which are asymmetric, and for the outliers at p≤0.001. In **Fig. 2B&D** we compare the observed χ^2^ frequency distributions with the expected probability distribution. In this case again, the observed and expected χ^2^ distribution are indistinguishable. In **Fig. 2E**, we tabulate the observed probabilities from the simulations. We note that any errors in implementation of the method, such as incorrectly determined dF values, lead to deviations from the expected values in these simulations.

**Figure 2.**
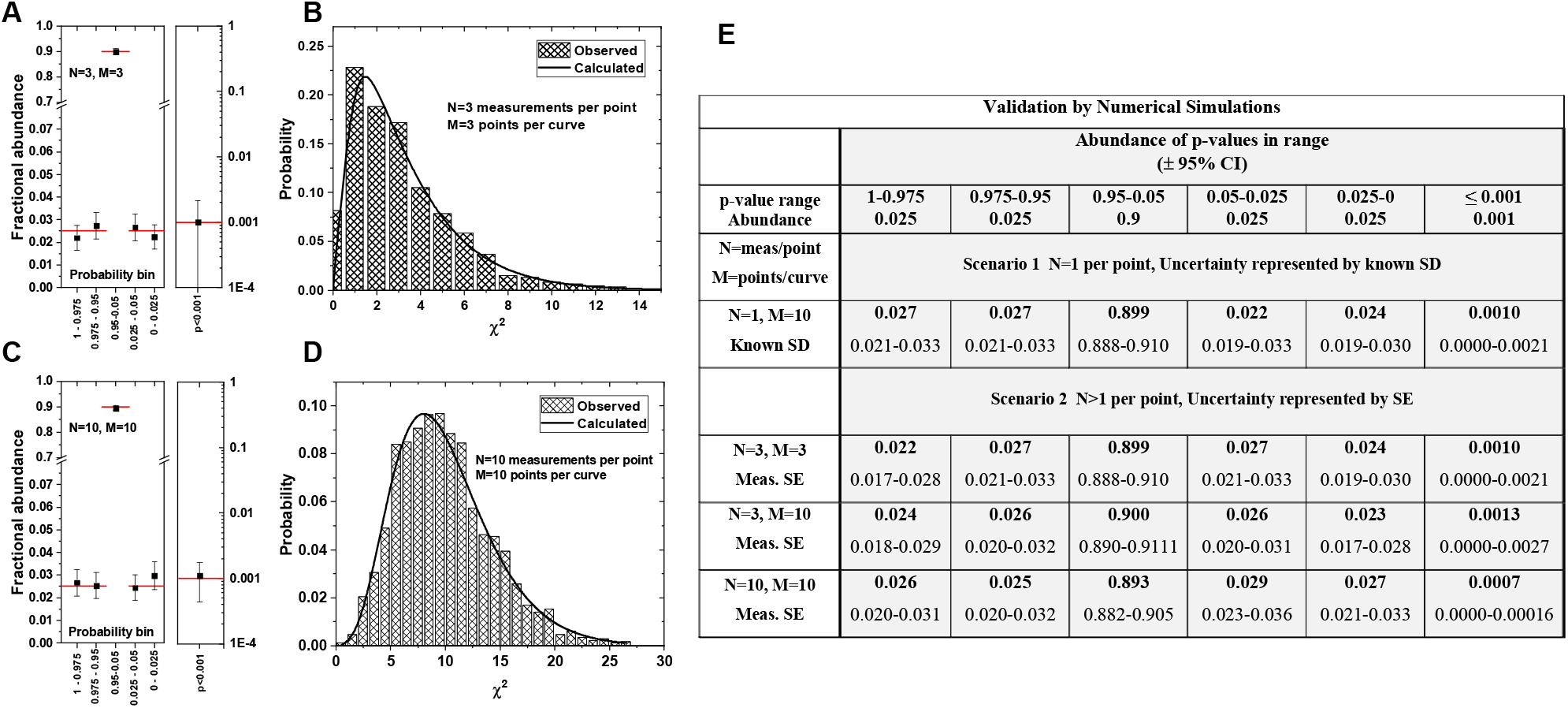
Validation of the modified Chi-squared method using numerical simulations. Pairs of curves were simulated according to both scenarios, N=1 with known SD and N>1 with calculated SE. We sampled M pairs of points where each point is comprised of 1 (scenario 1) or N (scenario 2) random samples from a Gaussian parent population. At least 6,000 simulated pairs of curves were compared. **A**,**B:** Simulations with N=3 and M=3. **C**,**D:** Simulations with N=10 and M=10. **A**,**C:** Expected and measured probability of calculated overall p-values, which are shown ± 95% CI. Horizonal bars indicate the expected values for each probability bin. The right box in each panel shows the data for p<0.001 on a log scale. **B**,**D:** Expected and measured distribution of overall χ^2^. **E**. The tabulated values show measured abundance of observed p-values for each range, for four examples, including the two scenarios described here. For each, a 95% confidence interval is also listed. All observed abundances fall within the expected range.

### Post hoc analysis

The p-values determined for pairs of points calculated at each value of X using the t-test (Col P, above) enable the *post hoc* determinization of which pairs of points contribute most to the overall difference between the curves. To state a standard statistical significance for these multiple t-tests, one has to correct for multiple comparisons. For simplicity and stringency, we use the Bonferroni correction which enables calculation of a corrected cutoff for statistical significance. Formally, the Bonferroni correction can be described as:

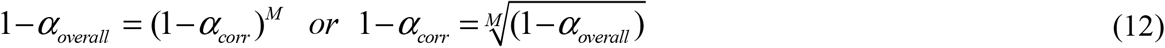

where α_overall_ is the desired total type 1 error rate over all M comparisons, typically 0.05, and α_corr_ is the corrected cutoff for significance of each of M comparisons needed to achieve the desired α_overall_. However, the Bonferroni-corrected α can be closely approximated by:

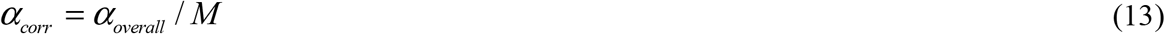

where the corrected cutoff for significance is equal to the uncorrected cutoff (0.05) divided by M, the number of pairs of points being compared. The power of the χ^2^ method is that a highly significant difference between curves can be identified even when none of the differences between the individual points rise to the level of significance.

We note that the Bonferroni correction may be considered inappropriately stringent for curves containing many comparisons. In this case, the Benjamini-Hochberg [4] or similar methods for performing multiple comparisons using a false discovery rate approach may be more appropriate.

### Multiple pairwise comparisons of curves

If more than one pairwise comparison between curves is needed, such as multiple curves versus a control or multiple curves versus each other, the χ^2^ method remains applicable, but a correction must be made for multiple comparisons between curves. Here again, the Bonferroni correction to α, the cutoff for statistical significance, is appropriate and statistically robust,

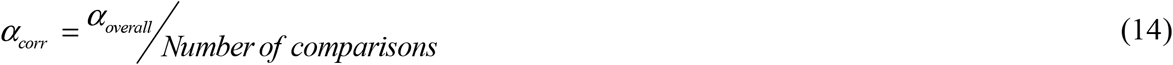

### Example data analysis

Next, we show the application of the Chi-squared method using a previously published data set [5] describing the heterodimerization of mutants of the receptor FGFR3. We characterized the heterodimerization of the receptor tyrosine kinase FGFR3 with two pathogenic mutants that cause skeletal and cranial abnormalities in humans, A391E and G380R. Experimentally, we measured autophosphorylation of the receptors as functions of ligand concentration, both in the presence and absence of enzymatically inactive truncated FGFR3, called ECTM_WT_-1. If heterodimerization between WT and mutant occurs, the presence of ECTM_WT_-1 will inhibit the mutant receptor phosphorylation.

In **Fig. 3**, we show the phosphorylation versus ligand concentration for WT, A391E and G380R, with and without the inhibitory construct. Each point is comprised of three independent measurements of phosphorylation, and is shown as mean ± SEM.

**Figure 3.**
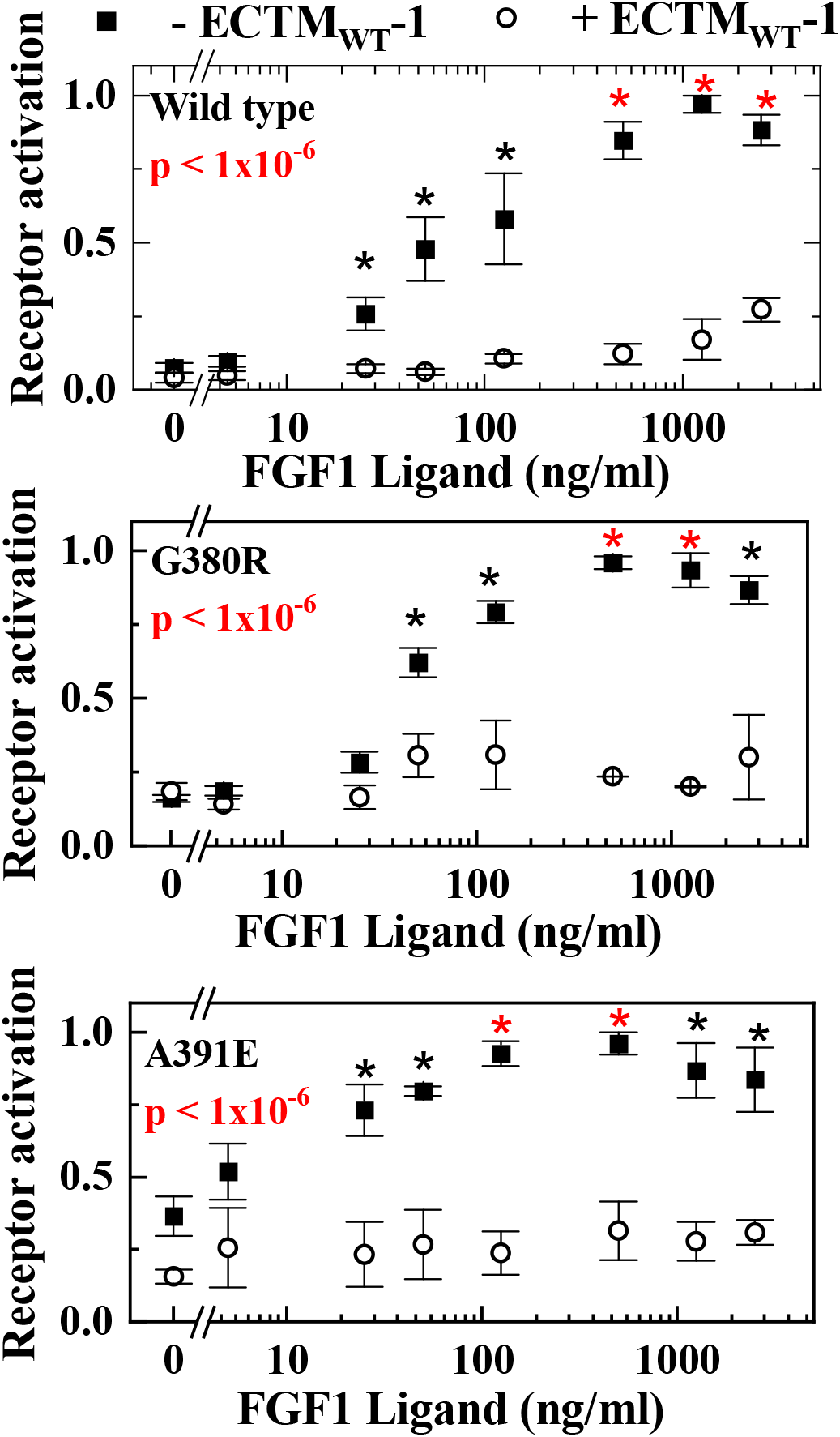
Curve-vs-curve comparisons of the phosphorylation of the kinase domain of the receptor FGFR3. As described by us previously [5], the activation levels for wild type and two mutants were measured in the presence and absence of an inhibitory wild type construct, ECTMwt-1, comprised of extracellular and transmembrane domains, but without a functional kinase domain. For each point, three independent experiments were done and values shown are means ± SE. In each case, comparison of the curves yielded p-values < 1x10^-6^. For secondary, post hoc analysis, we also added point-by-point comparisons wherein a black asterisk indicates a pair of points with p<0.05. A red asterisk indicates a pair of points that are significant after the Bonferroni correction for multiple comparisons, p<0.006.

We applied the modified χ^2^ method, described above, to the three pairs of curves. All pairwise comparisons are highly significant and give overall p-values of less than 1x10^-6^. The significance of each individual pair of points is calculated using a 2-sample t-test as described. Differences, with p<0.05 are shown with a black asterisk and those with p<α_corr_, the Bonferroni corrected cutoff of 0.0063 for the eight comparisons, are shown with a red asterisk.

When we compared the degree to which each sequence, WT, A391E and G380R, is inhibited by the enzymatically inactive construct, the curves are superficially similar as shown in **Fig. 4A**. The Chi-squared method we describe here was initially developed to address whether these curves are significantly different [5], however the correction for the deviation from normality was not applied at that time. Here, we reanalyze the three comparisons with the fully developed Chi-squared method.

**Figure 4.**
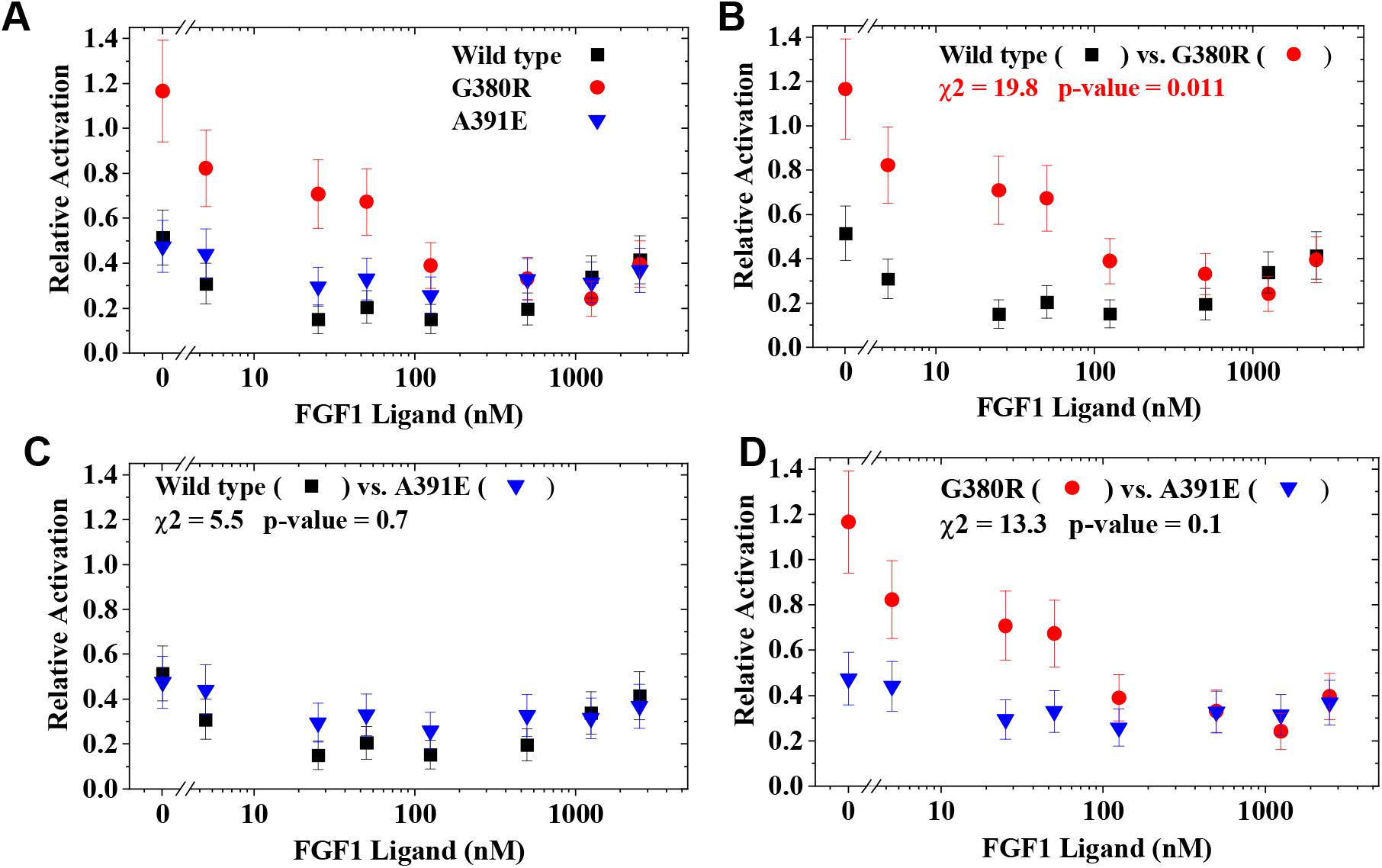
Curve-by-curve comparisons of the relative activation calculated by taking the ratio of receptor activation with/without ECTMwt from the previously published experiments [5], shown in **Fig. 3**. A value of 1 indicates no inhibition by ECTMwt and a value of 0 indicates complete inhibition. For each point, three independent ratios14were averaged. Values shown are means and error bars are SE. Chi-squared values and p-values for the curve-by-curve comparisons are given in panels, **B, C** and **D**.

All points in **Fig. 4** are the result of 3 independently measured ratios of activation with and without the inactive construct. Means and standard errors are shown. There are 8 pairs of points in each comparison. The three activation curves are shown together in **Fig. 4A** and the pairwise comparisons and modified Chi-squared method results are shown in **Fig. 4B-D**. Because we are making three pairwise comparisons, we apply the Bonferroni correction to obtain an adjusted cutoff of α=0.05/3 = 0.017, for statistical significance of the overall p-value. Based on this method, we conclude that there is statistically significant difference between the inhibition of wild type versus the G380R mutant, that is only slightly smaller than the cutoff for significance, and that there is no significant difference in the other comparisons.

In **Table 1** we provide the relative activation data that is plotted in panel **Fig. 4B** for wild type FGFR3 and the G380R mutant. In **Table 2**, we provide the intermediate Chi-squared calculations outlined above. These example data and calculation results are provided for users to validate any spreadsheet or coded application of the modified Chi-squared method.

**Table 1.**
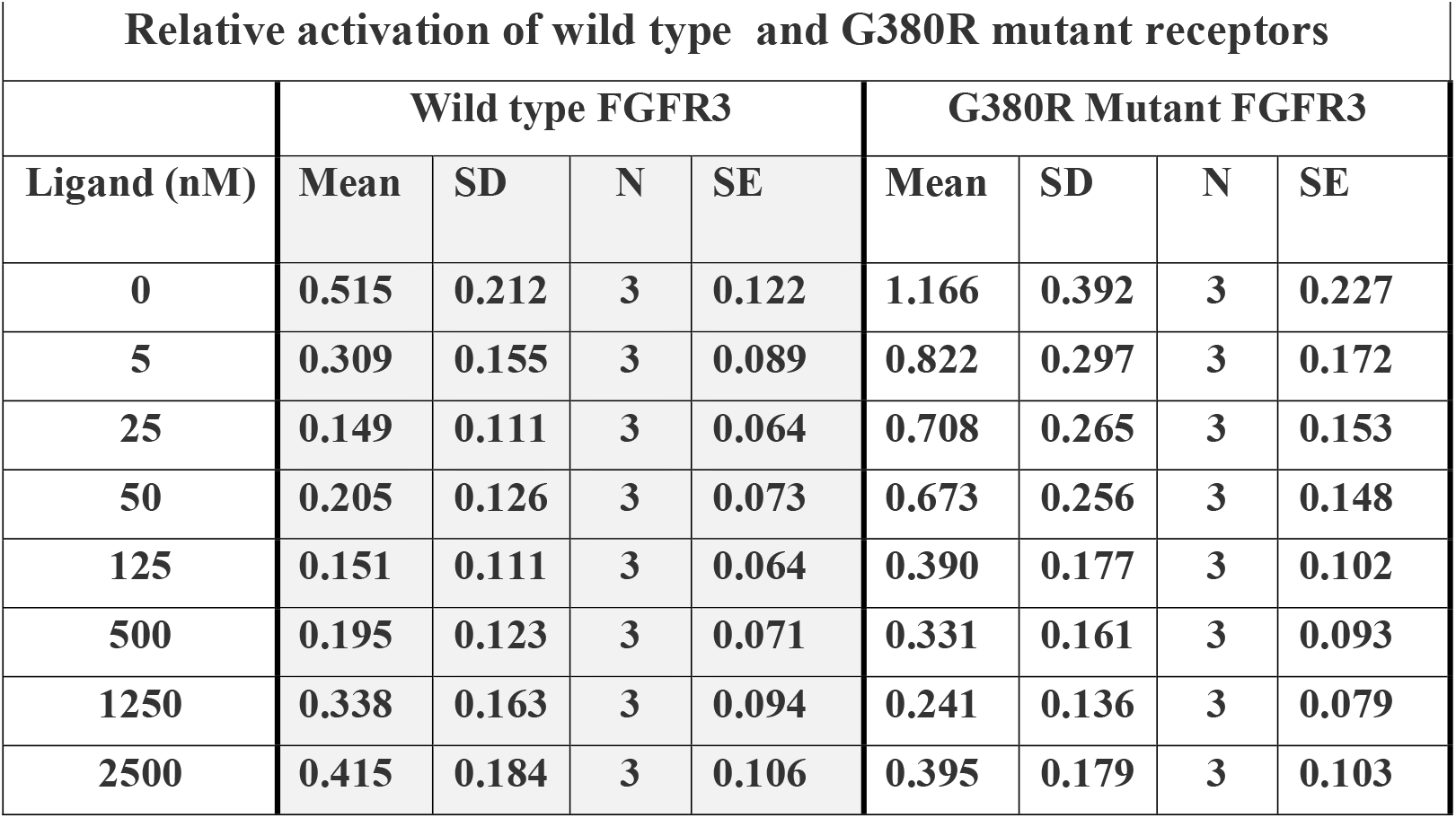
Example data from the analysis shown in **Fig 4B**. Mean relative activation data [5] are given here for the pair of curves shown in **Fig. 4B**. At each ligand concentration, we measured relative activation in 3 independent experiments. The data in the table are the mean, SD, N and SE for each combination of mutant and ligand concentration.

**Table 2.** Example calculations of the Chi-squared test applied to the two curves shown in **Fig. 4B** and in **Table 1**. The values in the table are the intermediate calculations in the Chi-squared test and the final result. In this case the sum of Chi-squared is 19.8, which at 8 degrees of freedom is equivalent to a p-value of 0.011.

| **$\chi^2$ based comparison of wild type versus G380R mutant curves** |  |  |  |  |  |  |  |
| --- | --- | --- | --- | --- | --- | --- | --- |
| **Ligand (nM)** | **Difference** | **SE Diff** | **t-value** | **N/pt** | **dF** | **P-value** | **$\chi^2$** |
| **0** | **0.651** | **0.257** | **2.527** | **3** | **4** | **0.065** | **3.409** |
| **5** | **0.513** | **0.194** | **2.652** | **3** | **4** | **0.057** | **3.627** |
| **25** | **0.558** | **0.166** | **3.363** | **3** | **4** | **0.028** | **4.814** |
| **50** | **0.468** | **0.165** | **2.841** | **3** | **4** | **0.047** | **3.952** |
| **125** | **0.238** | **0.121** | **1.972** | **3** | **4** | **0.12** | **2.419** |
| **500** | **0.136** | **0.117** | **1.161** | **3** | **4** | **0.31** | **1.029** |
| **1250** | **0.097** | **0.123** | **0.791** | **3** | **4** | **0.47** | **0.514** |
| **2500** | **0.020** | **0.148** | **0.135** | **3** | **4** | **0.90** | **0.016** |
|  |  |  |  |  | **Sum of $\chi^2$** |  | **19.8** |
|  |  |  |  |  | **dF** |  | **8** |
|  |  |  |  |  | **p-value** |  | **0.011** |

### Restrictions and assumptions

A limitation of this modified Chi-squared method for comparing curves is that the two curves must have data measured at the same values of X to enable pairwise comparisons. Data from one curve without corresponding data at the same X in the other must be ignored, or an appropriate extrapolation of data and uncertainty must be carried out using nearby points. With large data sets containing many points, binning data is an effective way to ensure that curves can be compared at the same X-values. Bins do not have to be equally spaced, or equally populated, but binning can only be done over ranges of X for which the Y value does not change substantially.

A second significant restriction is that the data, at each value of X, must approach a Gaussian distribution for large N, an assumption that is usually satisfied in biological data sets. Dataset normality can be tested with a D-Agostino-Pearson [6] or other normality test. Non-Gaussian data can be log-transformed, which frequently gives a normal distribution, when the linear data are not Gaussian. Use of log transformed data often provides a rigorous solution to the problem of non-normal data.

A third restriction is that SD/SE values cannot be zero for both curves to be compared. This problem can arise in experiments with a threshold concentration or dilution number describing a binary outcome, for example in bacterial growth inhibition assays performed via serial dilution. In this case one must assume a minimal nonzero value of SD/SE, which is based on an average uncertainty across many such measurements.

## Conclusions

We present here a simple modified Chi-squared based method to determine the statistical significance of the difference between two arbitrary curves. This simple and rigorous method can be performed using a spreadsheet program and is applicable to any pair of X-Y datasets in which the X-values are matched in pairs and in which the uncertainties of the individual points are known or have been measured in independent experiments. A functional Excel spreadsheet is provided as **Supplemental Information**.

## Supporting information

Excel Spreadsheet for Chi squared calculation

